# Near Infrared Nanosensors Enable Optical Imaging of Oxytocin with Selectivity over Vasopressin in Acute Mouse Brain Slices

**DOI:** 10.1101/2022.10.05.511026

**Authors:** Nicole Navarro, Sanghwa Jeong, Nicholas Ouassil, Jaewan Mun, Esther Leem, Markita P. Landry

## Abstract

Oxytocin plays a critical role in regulating social behaviors, yet our understanding of its role in both neurological health and disease remains incomplete. Real-time oxytocin imaging probes with the spatiotemporal resolution relevant to its endogenous signaling are required to fully elucidate oxytocin function in the brain. Herein we describe a near-infrared oxytocin nanosensor (nIROx), a synthetic probe capable of imaging oxytocin in the brain without interference from its structural analogue, vasopressin. nIROx leverages the inherent tissue-transparent fluorescence of single-walled carbon nanotubes (SWCNT) and the molecular recognition capacity of an oxytocin receptor peptide fragment (OXTp) to selectively and reversibly image oxytocin. We employ these nanosensors to monitor electrically stimulated oxytocin release in brain tissue, revealing oxytocin release sites with a median size of 3 μm which putatively represents the spatial diffusion of oxytocin from its point of release. These data demonstrate that covalent SWCNT constructs such as nIROx are powerful optical tools that can be leveraged to measure neuropeptide release in brain tissue.

## Introduction

Oxytocin is a nonapeptide^1^ that plays essential roles in mammalian social and reproductive behavior^2^. Synthesized predominantly in the hypothalamus, oxytocin acts both peripherally as a peptide hormone and centrally as a neuropeptide to perform distinct but complementary functions^3^. When released throughout the brain, neuropeptide oxytocin regulates complex emotions and social behaviors^4^, including recognition^5,6^, bonding^7^, and anxiolysis^8,9^. Oxytocin is also implicated in the pathogenesis of various social impairment disorders such as generalized social anxiety disorder (GSAD)^10,11^ and autism spectrum disorder (ASD)^12,13^ and has thus garnered interest as a potential therapy and therapeutic target.

While neuropeptide oxytocin has been studied extensively, we lack the tools to directly probe oxytocin signaling at the spatial (μM) and temporal (s) scales at which it is hypothesized to signal, precluding elucidation of its function in oxytocinergic communication. The gold standard for central oxytocin detection involves sampling with microdialysis followed by quantification with radioimmunoassay (RIA) or enzyme-linked immunosorbent assays (ELISA). The limitations of microdialysis include poor spatiotemporal resolution and significant variability in reported oxytocin concentration with different sample processing methods and quantification assays^14–16^. While fast-scan cyclic voltammetry (FSCV) can be used to directly measure neurotransmitter release with nanomolar sensitivity and high temporal resolution^17^, FSCV-based detection of neuropeptides has heretofore proved challenging. Einaga *et al.* developed a boron-doped diamond electrode for oxytocin detection^18^, but the method remains limited to use in expert laboratories.

More recently, Kwon and colleagues developed a genetically-encoded oxytocin sensor, OXTR-iTango2, which enables EGFP expression in the presence of both oxytocin and blue light^19^. Utilizing the endogenous oxytocin receptor (OXTR)^20^ as a sensing moiety, which binds both oxytocin and vasopressin, OXTR-iTango cannot distinguish between these two structurally analogous neuropeptides. This optogenetic platform enables labeling of oxytocin sensitive neurons for post-experimentation analysis of oxytocinergic cell activation but lacks the temporal resolution requisite for real-time oxytocin imaging. Selectivity, particularly against vasopressin, and slow temporal resolution limitations continue to pose a challenge for OXTR- and expression-dependent sensors, respectively^21^.

The selectivity and spatiotemporal limitations of current techniques motivate this study, whereby we employ single-walled carbon nanotube (SWCNT)-based probes to image oxytocin in the brain. SWCNT are inherently fluorescent in near-infrared tissue-transparent wavelengths, biocompatible, photostable, and functionalizable with biopolymers to impart molecular recognition towards targets of interest^22,23^. In prior work using SWCNT to image neuromodulators, Beyene *et al.* developed a catecholamine nanosensor non-covalently functionalized with short oligonucleotides to image dopamine release both in striatal brain tissue^24,25^ and at the level of single dopamine release sites in neuronal soma and dendrites^26^.

Until recently, covalent modification of the SWCNT surface for nanosensor development remained intractable. Such covalent modifications normally introduce defects that quench SWCNT fluorescence, however, Setaro and colleagues developed a strategy to enable direct covalent functionalization of SWCNT while retaining the *sp^2^* carbon network and ergo their fluorescence^27,28^. As described herein, we leveraged this strategy and the recognition capacity of an oxytocin receptor peptide fragment (OXTp)^29^ to develop realtime oxytocin imaging probes.

### Oxytocin nanosensor development and *in vitro validation*

Near-infrared oxytocin nanosensors (nIROx) were prepared from SWCNT as follows: (1) chemical re-aromatization of pristine SWCNT followed by conjugation with glycine to generate carboxylated triazine SWCNT (COOH-Trz-SWCNT), (2) covalent attachment of the oxytocin receptor peptide fragment (OXTp) via amide bond formation to make OXTp-modified SWCNT (OXTp-SWCNT), and (3) noncovalent adsorption of poly-cytosine (C_12_) to produce nIROx (Fig. 1a). Defect-free fluorescent Trz-SWCNT were generated from pristine HiPCo SWCNT and subsequently reacted with glycine to form COOH-Trz-SWCNT by following previously reported protocols^27,28,30^. This synthetic strategy does not disrupt the *sp^2^* carbon network of SWCNT, thus retaining their near-infrared fluorescence. COOH-Trz-SWCNT were conjugated to the oxytocin peptide by amide bond formation between the N-terminal amine of the oxytocin peptide and the carboxyl group in COOH-Trz-SWCNT. The oxytocin peptide contains both a spacer region (GPGSG) at the N-terminus and a peptide sequence derived from the transmembrane VI domain of the human oxytocin receptor (MTFIIVLAFIVCWTPFFFV). Previous work demonstrates that the oxytocin peptide binds to oxytocin with high affinity^29^. Attachment of the oxytocin peptide to the SWCNT surface was verified through X-ray photoemission spectroscopy (XPS). The sulfur 2p/carbon 1s ratio increased from 0.05 to 0.12 after oxytocin peptide attachment to the carboxylated triazine ring due to the cysteine residue in the oxytocin peptide, while the peak area ratio of nitrogen 1s/carbon 1s was unchanged (Supplementary Figure 1). To impart colloidal stability to OXTp-SWCNT, we leveraged a previously established protocol to noncovalently pin C_12_ DNA to the SWCNT surface. The C_12_ sequence was selected for its stable noncovalent adsorption to the SWCNT surface and low affinity for oxytocin (Fig. 1e)^31^.

**Fig. 1.**
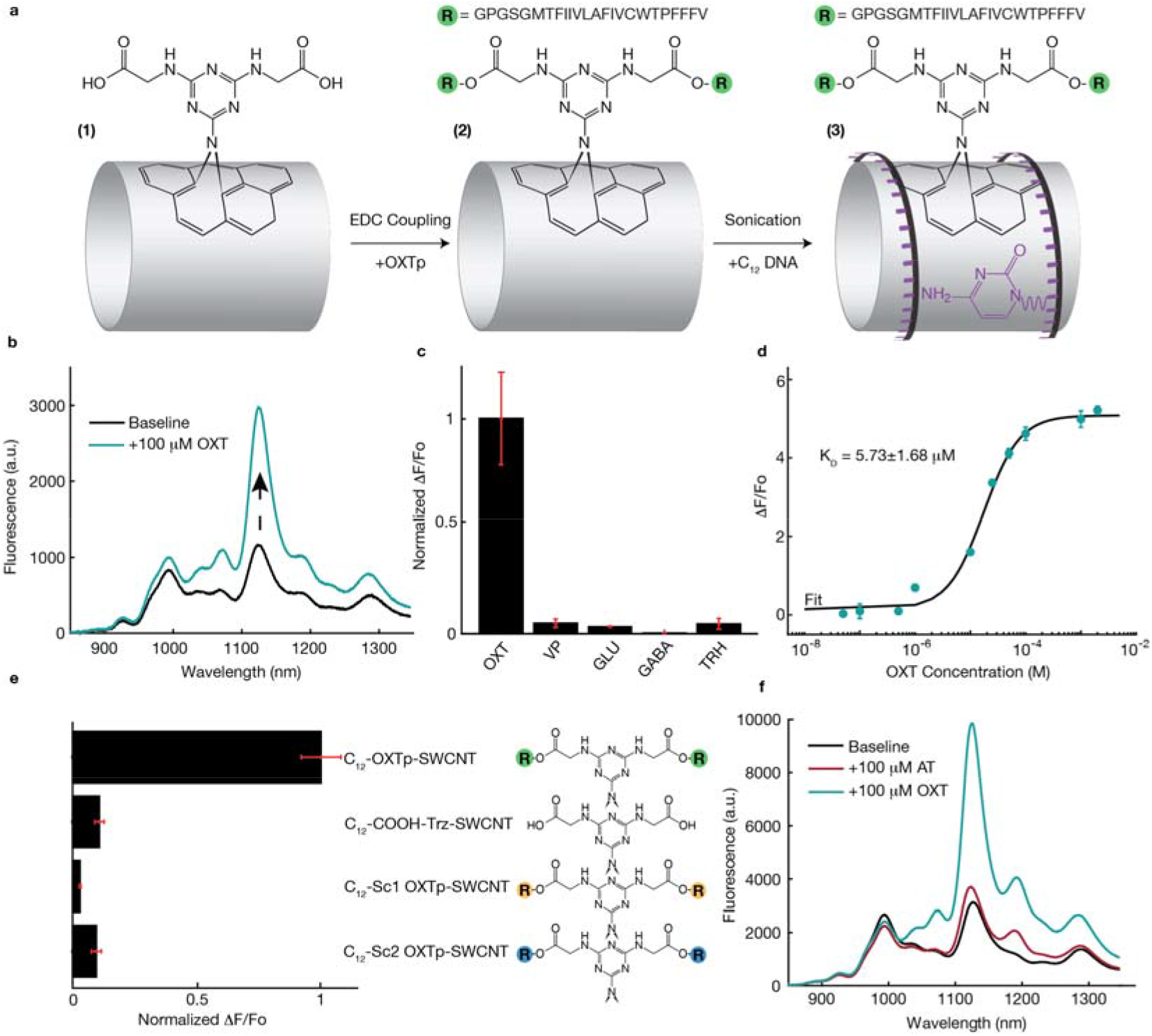
nIROx synthesis and *in vitro* validation. **a,** Synthesis scheme of nIROx. The first product, COOH-Trz-SWCNT, was synthesized through covalent attachment of triazine to pristine SWCNT followed by glycine conjugation. The oxytocin receptor peptide (OXTp) was subsequently attached via EDC coupling. OXTp-SWCNT were noncovalently functionalized with C_12_ DNA to form colloidally stable nIROx. **b,** The full fluorescence spectrum of nIROx before (black) and after (blue) the addition of 100 μM oxytocin (OXT) in PBS. The maximum turn-on fluorescence response is observed at the center wavelength of the (9,4) SWCNT chirality (~1126 nm). **c,** nIROx selectivity screening at 50 μM oxytocin (OXT), vasopressin (VP), glutamate (GLU), *γ*-aminobutyric acid (GABA), and thyrotropin-releasing hormone (TRH). Black bars represent the mean ΔF/F_o_ at 1126 nm from n=3 replicates normalized to OXT response, with standard deviation of these replicates shown in red. **d,** Dose response curves of nIROx for OXT. Blue circles and error bars represent the mean of n=3 experimental measurements and the standard deviation of these measurements, respectively. ΔF/F_o_ is calculated from the normalized change in peak intensity at 1126 nm. The black line represents the cooperative binding model fit to experimental data. The Kd value is reported with 95% confidence intervals using the t-distribution. **e,** Response of various nanosensor constructs to 50 μM OXT, demonstrating that the oxytocin peptide sequence is requisite for OXT response and C_12_ DNA has low affinity for OXT. Black bars represent the mean ΔF/F_o_ at 1126 nm from n=3 replicates normalized to nIROx response, with standard deviation of these replicates shown in red. **f,** Full fluorescence spectra of nIROx before (black) and after the addition of 100 μM atosiban (red), followed by the addition of 100 μM OXT (blue). Atosiban reduces nIROx response to OXT by ~30% *in vitro* by competitively inhibiting oxytocin analyte binding.

The resulting nanosensor, nIROx, exhibits a chirality-dependent fluorescence response to oxytocin, where the maximum turn-on response is observed at the center wavelength of the (9,4) SWCNT chirality (~1126 nm) (Fig. 1b). nIROx demonstrates a maximum peak fluorescence response (ΔF/F_o_) of 5.2 ± 0.10 (mean ± SD) to 2 mM oxytocin *in vitro* and responds to oxytocin in a concentration-dependent manner with a 235 nM limit of detection (LOD)^32^ (Fig. 1d, Supplementary Equation 1). We next calculated the nIROx kinetic parameters by fitting nIROx response versus oxytocin concentration to a cooperative binding model^33^, resulting in an equilibrium dissociation constant (K_d_) of 5.73 μM (Fig. 1d, Supplementary Equation 2). As positive controls, we constructed SWCNT-peptide conjugates from two scrambled oxytocin peptide sequences, and one SWCNT without peptide altogether (Fig. 1e). These negative controls, C_12_-COOH-Trz-SWCNT, C_12_-Scrambled OXTp 1-SWCNT, and C_12_-Scrambled OXTp 2-SWCNT responded to 50 μM oxytocin with normalized ΔF/F_o_ = 0.11 ± 0.02, 0.030 ± 0.004, and 0.10 ± 0.02 (mean ± SD), respectively, relative to nIROx. The insensitivity of these constructs to oxytocin confirms that nIROx response to oxytocin occurs via molecular recognition between the oxytocin peptide sensing moiety and oxytocin, and that this molecular recognition is perturbed when the peptide is scrambled or outright omitted, as expected.

To characterize nIROx selectivity, nanosensor response was evaluated for a panel of neurochemicals including: glutamate, *γ*-aminobutyric acid (GABA), vasopressin, and thyrotropin-releasing hormone (TRH), where nIROx was found to be insensitive to these analytes (Fig. 1c). As thyrotropin-releasing hormone is released in hypothalamus alongside oxytocin, nIROx selectivity to oxytocin over thyrotropin-releasing hormone [ΔF/F_o_ = 0.046 ± 0.025 (mean ± SD)] suggests that nIROx may be used as a selective oxytocin imaging probe in hypothalamic brain tissue ^34,35^. nIROx also exhibits selectivity for oxytocin over carbetocin [ΔF/F_o_ = −0.059 ± 0.021 (mean ± SD)] (Supplementary Figure 2a), a synthetic analogue with nanomolar binding affinity for the oxytocin receptor ^36^. Most notably, nIROx respond minimally to vasopressin with ΔF/F_o_ = 0.050 ± 0.019 (mean ± SD) relative to oxytocin. Vasopressin binds the endogenous oxytocin receptor with high affinity^37^ and differs from oxytocin by only 2 amino acid residues^38^. To date, our nanosensor is the first oxytocin imaging probe capable of distinguishing between these structurally analogous neuropeptides.

Prior to testing our probe in brain tissue, we evaluated nIROx reversibility by immobilizing nanosensors to the surface of a glass slide, enabling sequential oxytocin addition and removal from surface-adsorbed nIROx. We immobilized nIROx on a glass substrate using a previously described drop-casting method^33^ and tracked the integrated fluorescence of nIROx during multiple oxytocin washes. Nanosensor response was immediate upon addition of 100 μM oxytocin (Fig. 2a,b). Washing the glass surface with phosphate buffered saline (PBS) to remove oxytocin resulted in an immediate decrease in integrated fluorescence, while subsequent additions of 100 μM oxytocin restored nIROx response. Each nanosensor, represented by grey traces in Fig. 2a, maintained its fluorescence over continuous laser illumination over the course of the 300 second experiment without photobleaching. These data suggest that nIROx can reversibly bind and respond to oxytocin without signal attenuation.

**Fig. 2.**
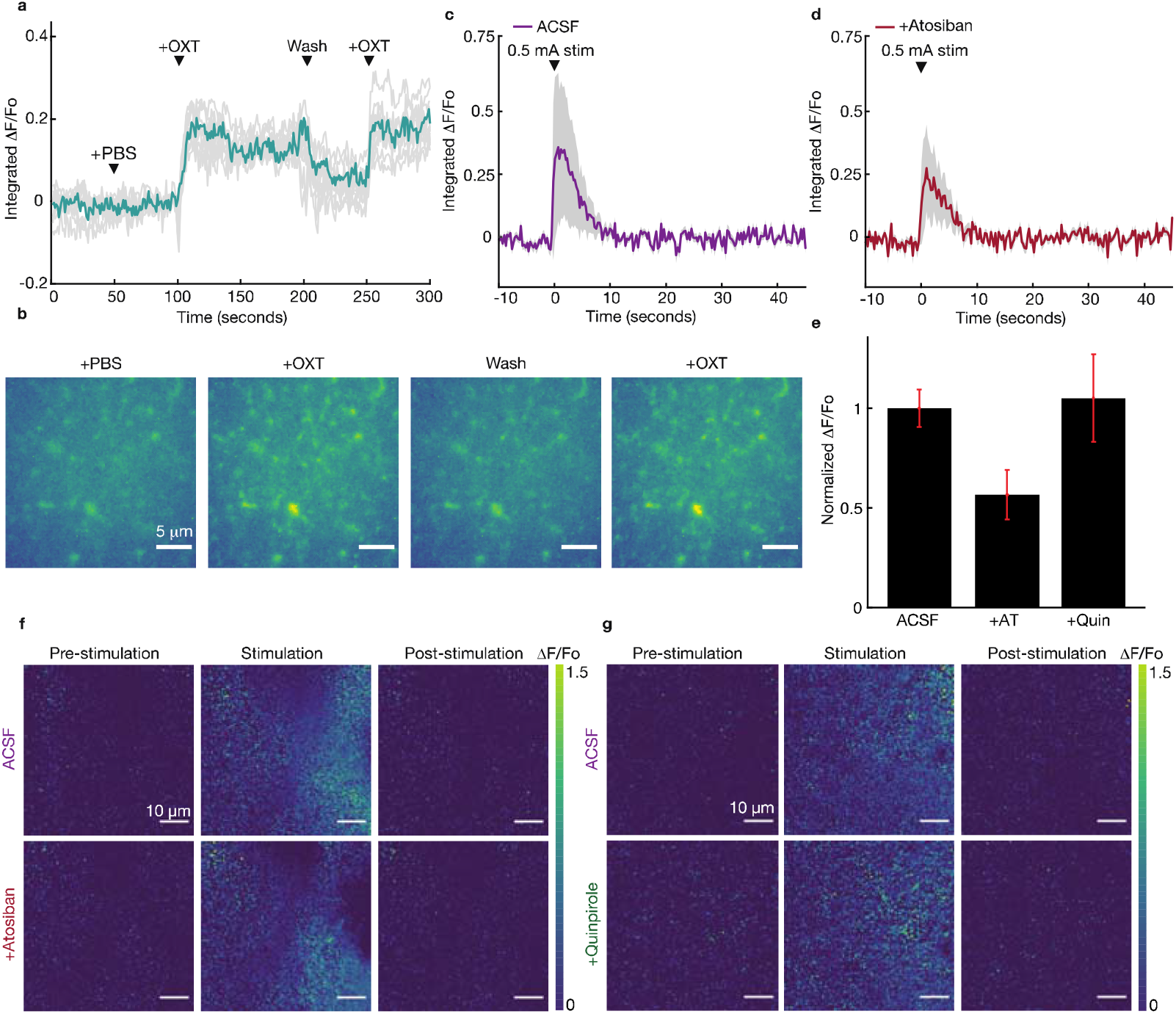
Reversible oxytocin imaging on solid substrates and in acute brain slices. **a**, *In vitro* integrated ΔF/F_o_ traces from single ROIs (gray) and the mean ΔF/F_o_ trace (blue) from two washes of 100 μM OXT on glass-immobilized nIROx (n = 12). nIROx nanosensor retains its sensitivity to oxytocin after substrate immobilization and demonstrates a reversible turn-on fluorescence response upon repeated oxytocin washes. **b,** *In vitro* three intensity heat maps within the same field of view from two washes of 100 μM OXT on glass-immobilized nIROx. Four frames are represented: “+PBS” after addition of PBS, “+OXT” after addition of 100 μM OXT, “Wash” after washing the glass with PBS, and “+OXT” after the second addition of 100 μM OXT. Scale bars represent 10 μm. **c,** *In brain slice* mean ΔF/F_o_ time trace (purple) and standard deviation (gray) following a single 0.5 mA electrical pulse stimulation in standard ACSF in the paraventricular nucleus (PVN). **d,** *In brain slice* mean ΔF/F_o_ time trace (red) and standard deviation (gray) following a single 0.5 mA electrical pulse stimulation in 1 μM atosiban in ACSF in the PVN. **e,** Integrated ΔF/F_o_ following 0.5 mA electrical pulse stimulations in ACSF, 1 μM atosiban (AT), and 1 μM quinpirole (Quin) normalized to ΔF/F_o_ in ACSF. Black bars represent the mean integrated ΔF/F_o_ across 3 stimulations with the standard deviations from these stimulations shown in red. **f,g** *In brain slice* ΔF/F_o_ of oxytocin nanosensor within the same field of view following 0.5 mA electrical stimulation in standard ACSF and either 1 μM atosiban (f) or 1 μM quinpirole (g) in the PVN. Three frames are represented for each: “pre-stimulation” is the baseline nanosensor fluorescence before electrical stimulation, “stimulation” is immediately following electrical stimulation, and “poststimulation” is after nanosensor fluorescence has returned to baseline. The white scale bars represent 10 μm.

### Oxytocin imaging in acute brain slices

We next verified that nIROx can reversibly image oxytocin release in *ex vivo* brain tissue. Acute coronal mouse brain slices were prepared as previously described^39^ and incubated with 2 mg/L nIROx in oxygen-saturated artificial cerebrospinal fluid (ACSF). Slices were then rinsed with ACSF to remove unlocalized nanosensor and transferred to an ACSF-perfused chamber for 15 minutes prior to imaging. This labeling protocol was previously demonstrated by Beyene *et al.* to enable even, widespread SWCNT labeling within brain tissue^24^ and shown by Godin *et al.* to enable SWCNT localization in the extracellular space (ECS)^40^ (Supplementary Figure 4). To image nIROx, we used a custom-built visible (400-725 nm) and near-infrared (750-1700 nm) microscope capable of serial imaging on the same detector. A 785-nm laser was employed to excite nanosensor fluorescence, while a mercury bulb was used for bright-field imaging. Imaging channels were selected via sliding mirror, and both near-infrared and brightfield images were collected serially with a Ninox VIS-SWIR broadband camera (Raptor Photonics) with the corresponding dichroic filters and a 60x water-dipping objective (Nikon).

To confirm nIROx nanosensors are capable of responding to oxytocin in brain tissue, nIROx was first introduced in the dorsal striatum of acute brain slices and nanosensor response was characterized upon addition of 100 μM of exogenous oxytocin. After collecting ~30 seconds of baseline nIROx fluorescence in slice, oxytocin was manually injected into the 3 mL recording chamber containing ACSF. Exogenous oxytocin injection yielded an increase in nanosensor fluorescence [ΔF/F_o_ = 0.36 ± 0.10 (mean ± SD)] and thus validated nanosensor utility for imaging oxytocin in *ex vivo* brain slices (Supplementary Figure 3a,b). Nanosensor response to exogenous oxytocin was not immediate and likely attributable to slow diffusion of the oxytocin injection through the recording chamber and into the acute slice bath. Washing of the chamber with ASCF gradually reduced nIROx fluorescence, demonstrating the reversibility of nanosensor binding to oxytocin in brain tissue and supported by our *in vitro* reversibility experiments.

nIROx were then introduced into the paraventricular nucleus (PVN) of mouse brain slices to evaluate their utility as endogenous oxytocin imaging probes. Oxytocin neurons are predominantly found in the supraoptic nucleus (SON), PVN, and accessory nuclei of the hypothalamus. Oxytocin is synthesized by magnocellular and parvocellular neurons, which differ in size, function, projection sites, and mode of oxytocin release^41^. The PVN was selected for oxytocin nanosensor imaging experiments owing to its distinct, identifiable structure and relatively high concentration of oxytocin, as reported previously in brain tissue by both microdialysis^42^ and mRNA expression^43^ studies. The imaging field of view of our custom microscope (178 μm by 142 μm) was capable of imaging multiple oxytocin release events within the PVN, where the somatic diameters of parvocellular and magnocellular neurons are roughly 15 μm and 25 μm, respectively^44^. To investigate nIROx imaging efficacy following electrically stimulated oxytocin release, we employed a bipolar stimulating electrode and applied a 0.5 mA, 1 millisecond single square pulse within the PVN for 2 biological replicates per experiment. Electrical stimulation evoked an instantaneous increase in nIROx fluorescence [ΔF/F_o_ = 0.41 ± 0.038 (means ± SD); *n* = 3] as shown in Fig. 2c and Supplementary Figure 8.

Upon demonstration that atosiban, a nonapeptide oxytocin analogue, decreases nanosensor response to oxytocin *in vitro* (Fig. 1f, Supplementary Figure 2b,c), we sought to recapitulate this effect in viable brain tissue as a positive control. Acute brain slices containing the PVN were labeled with nIROx and incubated in 1 μM atosiban for 15 minutes prior to repeating the above-mentioned stimulated oxytocin release imaging across 2 biological replicates. Incubation with atosiban, followed by electrical stimulation of acute tissue slices, yielded a lower nanosensor response relative to atosiban-free slices, with atosiban ΔF/F_o_ = 0.23 ± 0.05 (means ± SD) and a post-drug to pre-drug ΔF/F_o_ amplitude ratio of 0.56 ± 0.12 (means ± SD) across three stimulations (Fig. 2d-f, Supplementary Figure 5). These data are consistent with *in vitro* experiments which revealed a 28 ± 7.9% decrease in peak ΔF/F_o_ upon incubation with atosiban compared to the atosiban-free control (Supplementary Figure 2c). As atosiban attenuates nanosensor response similarly upon both exogenous oxytocin addition *in vitro* and electrical stimulation *ex vivo*, our results suggest nIROx response in brain tissue is largely attributable to endogenous oxytocin release. These data also provide insights into the mechanism by which atosiban reduces nIROx nanosensor response. Atosiban likely binds the oxytocin peptide sensing moiety of nIROx to competitively inhibit analyte binding and consequently reduces nanosensor response to oxytocin. While atosiban is known to antagonize the oxytocin receptor *in vivo*^45^, the oxytocin receptor is minimally expressed within the PVN^46^, and atosiban has not been reported to modulate oxytocin release within the hypothalamus. The effect of atosiban *ex vivo* is therefore likely due to competitive inhibition of nIROx rather than physiological changes in oxytocin release.

Concurrent monitoring of multiple neurotransmitters, neuromodulators, and neuropeptides, and their pharmacological agents often used in neurobiology research, is an important goal. To this end, coronal slices were also incubated in quinpirole, a D2 receptor agonist that inhibits presynaptic dopamine release^47,48^. Quinpirole treatment was used to verify that off-target pharmacological agents have a minimal effect on nIROx response, motivating potential future use of nIROx concurrently with other neuromodulator probes. Upon demonstrating that quinpirole treatment does not modulate nIROx fluorescence *in vitro* (Supplementary Figure 2a), slices were incubated in 1 μM quinpirole for 15 minutes prior to imaging. As expected, quinpirole had a negligible effect on nIROx ΔF/F_o_ across 2 biological replicates, yielding a post-drug to pre-drug ΔF/F_o_ amplitude ratio of 1.05 ± 0.220 (means ± SD) across three stimulations (Fig. 2g, Supplementary Figure 6,7). These results indicate that nIROx do not interact with off-target but broadly-utilized pharmacological agents such as quinpirole in brain tissue. Taken together, the results from atosiban and quinpirole treatment suggest that nIROx response in PVN-containing brain tissue slices is attributable to oxytocin release.

### Oxytocin release site analysis

Using image stacks from nIROx-labeled and PVN-containing brain tissue slices subject to electrical stimulation, we employed custom MATLAB software to identify oxytocin release sites, termed regions of interest (ROIs), where post-stimulation ΔF/F_o_ is high. Overlay of ROIs with pre-stimulation images show that ΔF/F_o_ hotspots are not correlated to nanosensor labeling density of brain tissue (Fig. 3a), as we had previously determined for nanosensors of this class. This observation suggests that ΔF/F_o_ hotspots correspond to greater oxytocin release rather than greater nanosensor localization and is consistent with the findings of Beyene *et al.* upon imaging catecholamine release with DNA-SWCNT nanosensors^24^. More recently, Bulumulla and colleagues utilized a SWCNT-based nanofilm to image dopamine release with synaptic resolution and demonstrated that ROIs represent individual neurochemical release sites with reported ROI size as a proxy for the spatial spread, or volume transmission element, of dopamine release^26^. We conducted similar analyses with nIROx brain slice imaging data and found that stimulations repeated in triplicate within the same field of view and across biological replicates yielded comparable putative oxytocin release site size distributions and a median ROI size of 3 μm (Fig. 3b). Based on the findings of Bulumulla *et al.*, ROIs in our data likely correspond to single oxytocin release sites in the PVN while ROI size represents the spatial diffusion of oxytocin from these release sites. We quantified this putative oxytocin release site density by calculating the number of release sites per unit of PVN brain tissue area. This analysis revealed a median of 3 putative oxytocin release sites per 1000 μm^2^ of brain tissue area.

**Fig. 3.**
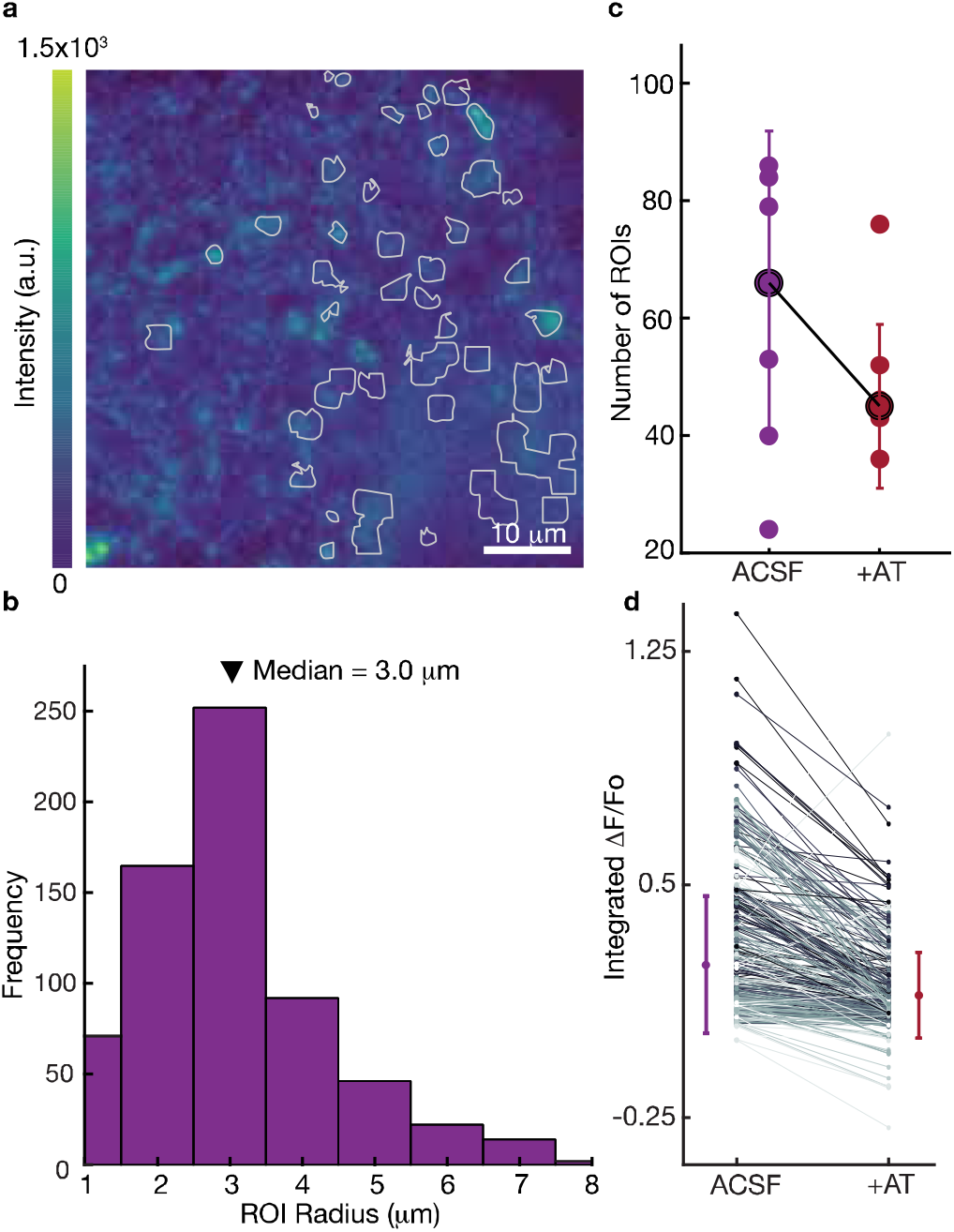
Oxytocin release site analysis of nIROx-labeled acute brain slices. **a,** A single frame from an image stack acquired in the paraventricular nucleus (PVN). The field of view is overlaid with ROIs (gray) identified programmatically by calculating ΔF/F_o_ time traces of the image stack. Color bar represents nanosensor labeling fluorescence intensity. Scale bar represents 10 μm. **b,** Frequency histogram of ROI sizes across 4 biological replicates and 12 stimulations of the PVN in ACSF with median ROI size of 3 μm. **c,** Distribution of the number of ROIs across 6 stimulations in ACSF (purple) and 1 μM AT in ASCF (red). Mean (black) and error bars representing standard deviation of these stimulations. **d,** Integrated ΔF/F_o_ and standard deviation for all 500 ROIs across all biological replicates in ACSF and 1 μM AT in ACSF where ΔF/F_o_ values represent experimental means across 3 stimulations.

While atosiban treatment did not significantly alter the ROI size distribution or median ROI size, as expected, atosiban did attenuate the number and release intensity of oxytocinergic ROIs. These results suggest that atosiban does not affect oxytocin diffusion from release sites for instances when oxytocin is successfully released, but rather interacts with nIROx to reduce nanosensor oxytocin sensitivity. Upon atosiban treatment, the post-drug field of view exhibits ~20% fewer ROIs relative to drug-free brain tissue stimulation. The total number of ROIs across all stimulations and biological replicates decreased from 366 to 297 following atosiban treatment, and the mean number of ROIs per stimulation decreased from 61± 25 to 49 ± 14 (means ± SD) with atosiban treatment (Fig. 3c, Supplementary Figure 9). These results suggest that atosiban “turns-off” a subpopulation of nIROx rendering these nanosensors non-responsive to oxytocin. We also compared ΔF/F_o_ values of ROIs before and after atosiban drug treatment. As the location and number of ROIs differs across stimulations, all 500 ROIs in the field of view were included in this analysis. This analysis revealed a post-to pre-drug ratio ΔF/F_o_ amplitude ratio of 0.75 ± 0.31 (means ± SD) across 3 stimulations (Fig. 3d). These analyses provide further insight into the mechanism by which atosiban affects nanosensor response, suggesting that atosiban reduces nIROx response to oxytocin in 2 ways: by both reducing the number of responsive nIROx nanosensors and simultaneously reducing the sensitivity of responsive nIROx nanosensors to oxytocin. These data in slice are consistent with *in vitro* nIROx reduction in sensitivity and support our hypothesis that nanosensor response in slice is attributable to oxytocin release.

## Conclusions

Imaging oxytocin in brain tissue with form factors compatible for use with existing neurobiological tools are needed to illuminate this missing dimension in mapping the chemical activity of oxytocinergic neural circuits. In particular, neuropeptides such as oxytocin are posited to influence broad networks of neuronal activity through diffusion beyond the synapse during neurotransmitter release and are hypothesized to play key roles in social and maternal behaviors. Importantly, oxytocin signaling disruption is a likely contributor to autism spectrum disorders based on studies observing amelioration in social behavior upon administration of oxytocin to autistic patients. However, tools to image oxytocin at spatial and temporal scales of relevance to its endogenous signaling, and with requisite selectivity over its vasopressin analog, remain non-existent.

Herein we describe the development and utilization of nIROx, an oxytocin imaging probe for use in acute brain tissue slices containing oxytocinergic brain regions. nIROx was synthesized through covalent attachment of an oxytocin receptor peptide fragment (OXTp) to pristine SWCNT using a strategy that maintains SWCNT near-infrared fluorescence. Our successful nIROx synthesis demonstrates that SWCNT re-aromatization chemistries can covalently incorporate molecular recognition elements on the SWCNT surface, a concept that could be extended to other neuropeptide, neurohormone, and neurotransmitter targets. The resulting nanosensor, nIROx, responds fluorescently to oxytocin in near-infrared wavelengths with a peak ΔF/F_o_ >500% *in vitro*. We further show that nIROx synthesized with scrambled OXTp sequences or without a peptide do not respond to oxytocin, confirming that our nanosensor response occurs through molecular recognition of oxytocin via the oxytocin peptide moiety. Response screens against physiologically relevant neurochemicals, such as thyrotropin releasing hormone (TRH) and oxytocin’s structural analogue, vasopressin, reveal nIROx selectivity for oxytocin. To date nIROx is the only oxytocin imaging probe developed that can distinguish between oxytocin and vasopressin for use in brain tissue.

Reversibility experiments both on solid substrates and in brain tissue revealed that nIROx responds reversibly to oxytocin and is capable of repeated imaging without signal attenuation and with no photobleaching. We also introduced our nIROx nanosensor into the PVN of acute brain slices and demonstrated that our nanosensor can image electrically stimulated release of oxytocin with an integrated ΔF/F_o_ of up to ~40%. From these imaging experiments, we identified ROIs, putative oxytocin release sites, measuring 3 μm in median diameter with a calculated density of 3 ROIs per 1000 μm^2^ in PVN tissue. Based on previous studies with SWCNT-based neuroimaging probes, it is likely that these ROIs likely represent oxytocin release sites while their size is an approximate measurement of oxytocin’s volume transmission upon release. To our knowledge, these data represent the first direct visualization of oxytocin in brain tissue with frame rate temporal and micron-scale spatial resolution. Lastly, we confirmed that nIROx nanosensor response *ex vivo* is insensitive to incubation with quinpirole, a D2 receptor agonist, but attenuated upon incubation with atosiban, a positive control that reduced nIROx response to oxytocin *in vitro*. We thus confirmed that nIROx response in brain tissue is attributable to oxytocin release and insensitive to off-target pharmacological agents. Importantly, the synthetic nature of nIROx and of all SWCNT-based probes potentiates their use in non-model organisms or young animals in which genetic manipulation is intractable or undesirable, which is of particular relevance for oxytocin given its presumed central role in early-age social development. Taken together, our results show that nIROx represents a nearinfrared fluorescent, non-photobleaching nanosensor selective for oxytocin imaging in acute tissue slices containing oxytocinergic regions of the brain.

## Acknowledgements

We acknowledge support of a Burroughs Wellcome Fund Career Award at the Scientific Interface (CASI) (MPL), a Dreyfus foundation award (MPL), the Philomathia foundation (MPL), an NIH MIRA award R35GM128922 (MPL), an NIH R21 NIDA award 1R03DA052810 (MPL), an NSF CAREER award 2046159 (MPL), an NSF CBET award 1733575 (to MPL), a CZI imaging award (MPL), a Sloan Foundation Award (MPL), a USDA BBT EAGER award (MPL), a Moore Foundation Award (MPL), and a DOE office of Science grant DE-SC0020366 (MPL). MPL is a Chan Zuckerberg Biohub investigator and a Hellen Wills Neuroscience Institute Investigator.

## Methods

### Materials

Small diameter HiPCo™ single-walled carbon nanotubes (SWCNT) were purchased from NanoIntegris (Batch #27-104). Oxytocin acetate salt hydrate, [Arg^8^]-vasopressin acetate salt, dopamine-HCl, thyrotropin-releasing hormone (TRH), glutamate, *γ*-aminobutyric acid, and atosiban were purchased from Millipore Sigma. Quinpirole and carbetocin were purchased from Tocris Bioscience. Peptide sequences were purchased from Thermo Fisher Scientific, and DNA sequences were purchased with standard desalting purification from Integrated DNA Technologies, Inc.

### nIROx synthesis

Nanosensors were fabricated by (1) the surface modification of SWCNT to form carboxylated SWCNT, (2) the conjugation of oxytocin binding peptide (OXTp) to carboxylated SWCNT to produce OXTp-conjugated SWCNT, and (3) DNA-wrapping of OXTp-conjugated SWCNT to form colloidally stable nIROx.

1. *Carboxylated SWCNT fabrication*: Fabrication of fluorescent carboxylated SWCNT was adapted from previous literature^27,28^. In brief, SWCNT (1 g) were dispersed in N methyl 2 pyrrolidone (150 mL) in a round bottom flask and bath sonicated (Branson Ultrasonic 1800) for 1 h. The dispersion was stirred for 1 h at 25 °C and cooled to 0 °C. 2,4,6 1,3,5 trichloro triazine (10 g, 54 mmol) was dissolved in N methyl-2 pyrrolidone (50 mL) and slowly added to the SWCNT dispersion at 0°C. Sodium azide (1.76 g, 27 mmol) was added to the mixture and stirred for 2 h at 0°C followed by 12 h stirring at 70°C to yield triazine-functionalized SWCNT (Trz-SWCNT). The product was purified by centrifugation, washed with water and organic solvents (acetone, toluene, then chloroform), and lyophilized for storage and characterization. Trz SWCNT (100 mg) were dispersed in dimethylformamide (DMF) (50 mL) and bath sonicated for 15 min at room temperature. Next glycine (10 mg) and a 1.5 molar excess of triethylamine were added to the mixture and stirred at 65 °C for 2 days. The product was purified by centrifugation and sequentially washed with DMF (2 × 4 mL) and water (2 × 4 mL). The carboxylated SWCNT product was lyophilized for characterization and stored at room temperature.
2. *OXTp-conjugated SWCNT fabrication:* Carboxylated SWCNT (21.7 mg) were dispersed in DMF (0.5 mL) and bath sonicated for 15 min at room temperature. 1-Ethyl-3-(3-dimethylaminopropyl)carbodiimide (EDC) (1 μmol) and N-hydroxysuccinimide (NHS) (2 μmol) were added to the mixture and vortexed for 20 min. The mixture was centrifuged, the supernatant was discarded to remove excess EDC and NHS, and the precipitate was subsequently re-dispersed in DMF (4 mL). OXTp (GPGSGMTFIIVLAFIVCWTPFFFV) (100 nmol) and 1.5 molar excess of N,N-diisopropylethylamine (DIPEA) were added to the mixture solution and vortexed for 12 h at room temperature. For constructs prepared with scrambled OXTp sequences, either Scrambled OXTp 1 (GPGSGWLIFTVMCTIPAFVFFIFV) or Scrambled OXTp 2 (GPGSGITFILVMFFVWFVAICTPF) peptides were used instead of OXTp. The product was purified by centrifugation and subsequent re dispersion in washes of DMF (2 × 4 mL) followed by washes of water (2 × 4 mL). The OXTp-SWCNT product was lyophilized for characterization and stored at room temperature.
3. *DNA-wrapping of OXTp-conjugated SWCNT for oxytocin nanosensor:* OXTp-conjugated SWCNT (0.1 mg) was added to PBS (0.99 mL) and mixed with poly-cytosine (C_12_) (10 μL, 1 mM). The resulting mixture was first bath sonicated for 2 min and then probe-tip sonicated (Cole-Parmer Ultrasonic Processor, 3-mm tip in diameter) for 10 min at 5 W power in an ice bath. The suspension was centrifuged for 30 min at 16,100 g to precipitate non-dispersed SWCNT, and the supernatant containing colloidal C_12_-wrapped OXTp-SWCNT (nIROx) was collected. The concentration of the nIROx suspension was calculated by measuring absorbance at 632 nm (NanoDrop One, Thermo Scientific) with an extinction coefficient of ε = 0.036 (mg/L)^-1^ cm^-1 22^

### Optical characterization and analyte screening

Near-infrared fluorescence spectra were collected using a custom built spectrometer and microscope as described previously^49^. Measurements were obtained with a 20X objective on an inverted Zeiss microscope (Axio Observer.D1) coupled to a spectrometer (SCT 320, Princeton Instruments) and liquid nitrogen cooled InGaAs linear array detector (PyLoN-IR, Princeton Instruments). Nanosensor suspensions were excited with a 721 nm laser (OptoEngine LLC) inside a polypropylene 384 well-plate (Greiner Bio-One microplate).

For analyte screening, the baseline near-infrared fluorescence spectrum of each nanosensor-containing well was collected. Either PBS or analyte diluted in PBS was added, and post-analyte fluorescence spectra were collected at 10-minute time points until the maximum oxytocin fluorescence response was achieved (~60 minutes). Responses were calculated and reported as ΔF/F_o_ = (F-F_o_)/F_o_, where F_o_ is the peak fluorescence after PBS incubation, and F is the peak fluorescence after analyte incubation. The peak fluorescence corresponds to the (9,4) SWCNT chirality, which has a maximum nearinfrared fluorescence at ~1126 nm.

For selectivity screening, all analytes were added at a final concentration of 50 μM. During oxytocin screening of various nanosensor constructs, oxytocin was added at a final concentration of 50 μM. For dose response experiments, final oxytocin concentration ranged from 50 nM to 2 mM.

For drug screening, the baseline near-infrared fluorescence spectrum of nIROx was first collected. Atosiban was subsequently added and incubated for 10 minutes before collecting post-drug fluorescence spectra. Finally, oxytocin was added and incubated for 10 minutes before collecting post-neuropeptide fluorescence spectra. Responses were calculated and reported as ΔF/F_o_ = (F-F_o_)/F_o_, where F_o_ is the peak fluorescence after atosiban incubation, and F is the peak fluorescence after oxytocin incubation. Both atosiban and oxytocin were added at a final concentration of 100 μM.

For *in vitro* experiments, nIROx suspensions were prepared at 5 mg/L in PBS. All measurements were obtained in triplicate, and reported results include the means and standard deviations of these measurements. All nanosensor batches were validated for oxytocin response *in vitro* prior to brain tissue imaging.

For reversibility experiments on glass, nIROx were immobilized on MatTek glass-bottom microwell dishes (35 mm petri dish with 10 mm microwell). To immobilize nanosensors, the dish was washed twice with PBS (150 μL). Nanosensors (100 μL, 2.5 mg/L in PBS) were then added, incubated for 10 minutes, and removed. The adish was washed twice again with PBS (150 μL). Surface-immobilized nIROx were imaged on an epifluorescence microscope (100x oil immersion objective) and a Ninox VIS-SWIR 640 camera (Raptor) and excited with a 721 nm laser. For each imaging experiment, the z-plane was refocused and 120 μL PBS was added prior to recording. Image stacks were collected with a 950 ms exposure time and 1 Hz frame rate for 5 minutes. PBS was added at frame 50 and oxytocin was added a final concentration of 100 μM at frame 100. At frame 200, the dish was washed with PBS, and at frame 250, oxytocin was added again at a final concentration of 100 μM. Image stacks were processed in ImageJ by applying a median filter (0.5-pixel radius) and rolling ball background subtraction (300-pixel radius). ROIs were manually identified and characterized using the ROI analyzer tool. Responses were calculated and reported as ΔF/F_o_ = (F-F_o_)/F_o_, where F_o_ is the mean integrated fluorescence after PBS incubation, and F is the peak fluorescence at each timepoint after oxytocin addition. The means and standard deviation of all 12 ROI ΔF/F_o_ values are reported.

### Acute slice preparation and nanosensor labeling

The mice used for acute slice imaging experiments were male, 14-17 weeks old, C57BL/6 strain, and purchased from Jackson Laboratory. After weaning at postnatal day 21 (P21), mice were group-housed with nesting material on a 12:12 light cycle. All animal procedures were approved by the UC Berkeley Animal Care and Use Committee (ACUC). Acute brain slices were prepared using established protocols^39^. Mice were first anesthetized with an intraperitoneal injection of ketamine and xylazine, followed by transcardial perfusion with cutting buffer (119 mM NaCl, 26.2 mM NaHCO_3_, 2.5 mM KCl, 1mM NaH_2_PO_4_, 3.5 mM MgCl4, 10 mM glucose). The brain was subsequently extracted, and its connective tissues were trimmed. The brain was mounted on a vibratome cutting stage (Leica VT 1000) to prepare 300 μm thick coronal slices containing the putative paraventricular nucleus. Slices were incubated at 37°C for 30 minutes in oxygen saturated ACSF (119 mM NaCl, 26.2 mM NaHCO3, 2.5 mM KCl, 1mM NaH_2_PO_4_, 1.3 mM MgCl_2_, 10 mM glucose, 2 mM CaCl_2_) and then transferred to room temperature for 30 minutes. Slices were maintained at room temperature for all acute slice imaging experiments.

For nanosensor labeling, slices were transferred into a brain slice incubation chamber (Scientific Systems Design, Inc., AutoMate Scientific) containing 5 mL of oxygen-saturated ACSF. Slices were incubated with nanosensor at a final concentration of 2 mg/L SWCNT for 15 minutes. To remove unlocalized nanosensor from brain tissue, slices were rinsed for 5 seconds with oxygen-saturated ACSF through 3 wells of a 24-well plate. Rinsed slices were transferred to the recording chamber and perfused with ACSF for 15 minutes before imaging. All acute slice imaging experiments were performed at 32°C.

### Microscope construction and acute slice imaging

*Ex vivo* slice imaging experiments were conducted using an upright epifluorescent microscope (Olympus, Sutter Instruments) mounted onto a motorized stage. A 785-nm continuous-wave diode-pumped solid-state laser with a maximum power of 300 mW and a near TEM00 top hat beam profile (Opto Engine LLC) was used for nanosensor excitation. The beam was expanded to a diameter of ~1 cm using a Keplerian beam expander composed of two plano-convex lenses (*f* = 25 and 75 mm; AR coating B, Thorlabs). The beam was passed through a fluorescence filter cube [excitation: 800 nm shortpass (FESH0800), dichroic: 900 longpass (DMLP990R), and emission: 900 longpass (FELH0900); Thorlabs] to a 60× Apo objective (numerical aperture, 1.0; working distance, 2.8 mm; water dipping; high nIR transmission; Nikon CFI Apo 60XW nIR). Emission photons collected from the sample were passed through the filter cube, focused onto a two-dimensional InGaAs array detector [500 to 600 nm: 40% quantum efficiency (QE); 1000 to 1500 nm: >85% QE; Ninox 640, Raptor Photonics], and recorded with Micro-Manager Open Source Microscopy Software^50^.

### Exogenous oxytocin and electrical stimulation evoked oxytocin imaging with nIR microscopy

During exogenous oxytocin imaging, a total of 600 frames were acquired at nominally 4 frames per second in the dorsal striatum. After collecting 100 frames of baseline, 100 μM oxytocin was injected into the 3 mL recording chamber. Oxytocin was then removed from the slice with ACSF perfusion at frame 400.

To electrically stimulate oxytocin release, a bipolar stimulation electrode (MicroProbes for Life Science Stereotrodes Platinum/Iridium Standard Tip) was positioned within the putative paraventricular nucleus using a 4x objective (Olympus XLFluor 4x/340). The stimulation electrode was introduced into the top of the brain slice and an imaging field of view nominally 50 μm from the stimulation electrode was chosen using a 60x objective. A total of 600 frames were acquired at nominally 4 frames per second, where stimulations were applied after 200 frames of baseline. Each stimulation was repeated three times within a field of view, with 5 minutes between each stimulation. Stimulation pulses were applied for 1 millisecond at 0.5 mA. During drug screening, imaging experiments were first conducted in drug-free ACSF. Then using the same field of view, either quinpirole or atosiban was added to the imaging chamber through ACSF perfusion. The brain slice was incubated in 1 μM of either drug for 15 minutes before imaging. Drug screening imaging experiments were acquired in biological duplicate.

### Image processing and data analysis of nanosensor fluorescence response

Imaging movie files were processed using a custom MATLAB application (https://github.com/jtdbod/Nanosensor-Brain-Imaging). ROIs were identified by first applying a 25×25 pixel grid mask to the image stack. The median pixel intensity and a median filter convolution within each ROI was then calculated. Post-stimulation and exogenous oxytocin fluorescence responses were calculated as ΔF/F_o_ = (F-F_o_)/F_o_, where F_o_ is the average intensity for the first 5% of frames and F is the dynamic fluorescence intensity. Significant ΔF/F_o_ traces were identified by thresholding with Otsu’s methods to differentiate ROIs from the background. For stimulation experiments, decay time constants (*τ*) were computed for significant ROIs by fitting each ΔF/F_o_ trace to a first-order decay process. Latency to peak was calculated as *t*_peak_ - *t*_stim_, where *t*_peak_ is the time at which maximum fluorescence occurs, and *t*stim is time of stimulation. The ΔF/F_o_ traces where latency to peak is greater than 5 seconds were assumed to result from stimulation artifacts and were thus removed from analysis. The maximum ΔF/F_o_ of each significant ROI trace was identified, and the ΔF/F_o_ of a stimulation or exogenous oxytocin wash was reported as the median of these values. For stimulation experiments, ROI size was determined by first identifying F_max_ within a significant ROI. All pixels with an intensity value >50% of F_max_ were included within the boundaries of an ROI. ROIs with a radius <1 μm were removed.

