## Supplementary Information for "Near Infrared Nanosensors Enable Optical Imaging of Oxytocin with Selectivity over Vasopressin in Acute Mouse Brain Slices"

**Table of contents:**

Supplementary Equation 1. Limit of detection (LOD) calculation.

Supplementary Equation 2. Cooperative binding model fit.

Supplementary Figure 1. nIROx synthesis validation.

Supplementary Figure 2. nIROx *in vitro* drug screening.

Supplementary Figure 3. nIROx reversibility in brain tissue.

Supplementary Figure 4. Imaging oxytocin release evoked by electrical stimulation.

Supplementary Figure 5. Analysis of oxytocin release evoked by electrical stimulation and the effect of an oxytocin receptor antagonist in the paraventricular nucleus.

Supplementary Figure 6. Analysis of oxytocin release evoked by electrical stimulation and the effect of an D2/D3 receptor agonist in the paraventricular nucleus.

Supplementary Figure 7. Analysis of oxytocin release evoked by electrical stimulation and the effect of an D2/D3 receptor agonist in the paraventricular nucleus.

Supplementary Figure 8. Analysis of oxytocin release evoked by electrical stimulation.

Supplementary Figure 9. Region of interest (ROI) analysis of acute slice images.

**Supplementary Equation 1 | Limit of detection (LOD) calculation.** The LOD of nIROx for oxytocin is calculated as the lowest oxytocin concentration that is statistically different from the blank.

LOD = (mean ΔF/F_ο_)_blank_ + 3(STD ΔF/F_ο_)_blank_

For the concentrations of oxytocin tested, the LOD is calculated as ΔF/F_ο_ = 0.161 This ΔF/F_ο_ value corresponds to 235 nM oxytocin based on the cooperative binding model fit.

**Supplementary Equation 2 | Cooperative binding model fit.** Dose response data were fit to the Hill equation (cooperative binding model) to quantify nIROx nanosensor kinetic parameters.

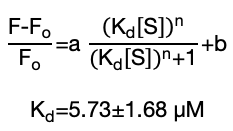

Kinetic parameters were calculated as n=1.33±0.51, b=0.14±0.29, a=4.94±0.5, and K_d_=5.73±1.68 µM and are reported with 95% confidence intervals using the t-distribution.

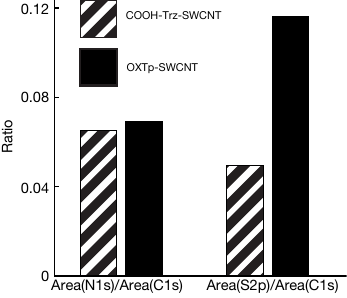

**Supplementary Figure 1 | nIROx nanosensor synthesis validation.** Verification of oxytocin peptide (OXTp) attachment through x-ray photoelectron spectroscopy (XPS). The sulfur 2p/carbon 1s ratio increases from 0.05 to 0.12 after OXTp conjugation due to the cysteine residue in the oxytocin peptide sequence. The nitrogen 1s/carbon 1s ratio remains unchanged.

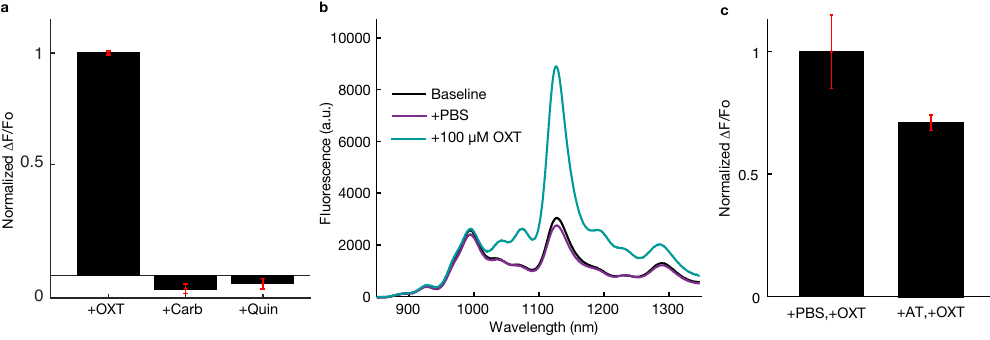

**Supplementary Figure 2 | nIROx *in vitro* pharmacological agent compatibility screening.**  **a,** nIROx optical response to 50 µM oxytocin (OXT), carbetocin (Carb), and quinpirole (Quin). Black bars represent the mean ΔF/F_ο_ at 1126 nm from n=3 replicates normalized to OXT response, with standard deviation of these replicates shown in red. **b,** Full fluorescence spectra of nIROx before (black) and after the addition PBS (purple), followed by the addition of 100 µM OXT (blue). **c,** nIROx optical response to 100 µM oxytocin after the addition of either 1X PBS or 100 µM atosiban (AT), and OXT receptor antagonist. Black bars represent the mean peak ΔF/F_ο_ at 1126 nm from n=3 replicates, and red error bars represent the standard deviation from these replicates. nIROx response to OXT decreases by 28% upon incubation with atosiban.

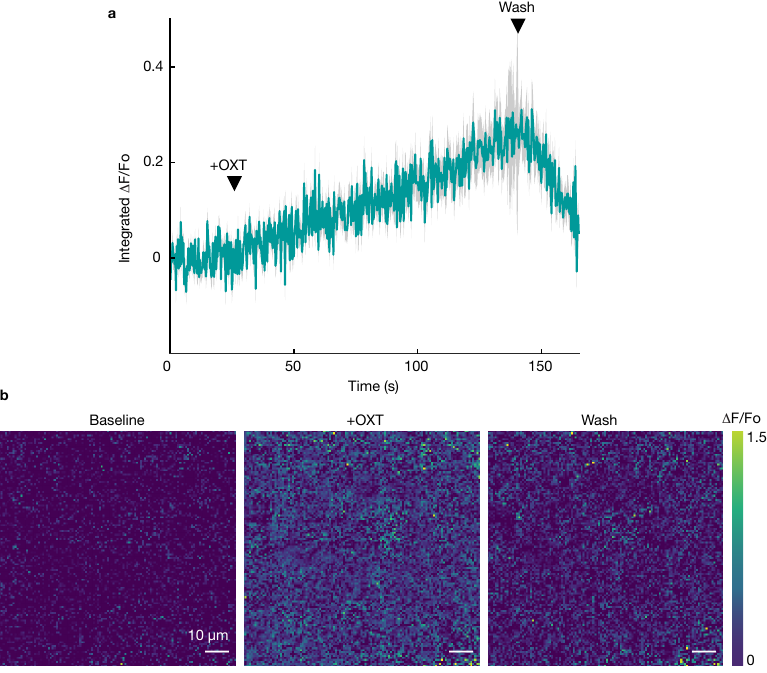

**Supplementary Figure 3 | nIROx nanosensor reversibility in brain tissue. a,** *In slice,* mean time trace (blue) and standard deviation (gray) of integrated ΔF/F_ο_ for nanosensors imbedded in an acute brain slice and exposed to an exogenous wash of 100 µM OXT. OXT was manually injected after 100 frames of baseline, and the slice was washed with an ACSF perfusion after 400 frames. nIROx sensitivity and reversibility upon immobilization is recapitulated in brain tissue. **b,** Three images within the same field of view of integrated ΔF/F_ο_ of nIROx in the dorsal striatum. Three frames are represented: “baseline” is before the addition of oxytocin, “+OXT” after the addition of 100 µM exogenous oxytocin, and “wash” is after washing the slice with ACSF to remove oxytocin. Scale bars represent 10 µm.

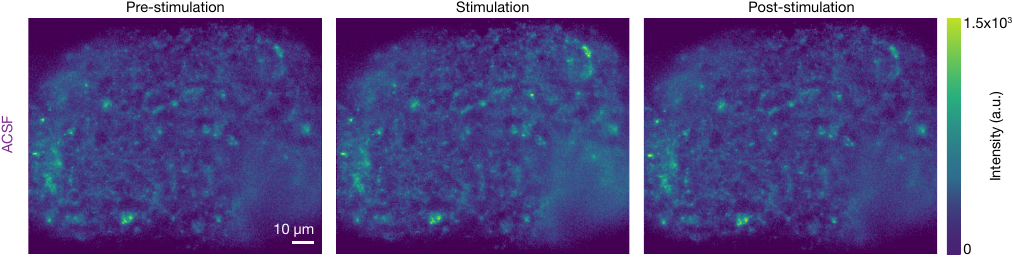

**Supplementary Figure 4 | Imaging oxytocin release evoked by electrical stimulation.** *In slice,* three images within the same field of view of nIROx-labeled acute brain slices following a 0.5 mA single-pulse electrical stimulation in standard ACSF in the PVN. “Pre” is the baseline before electrical stimulation, “stim” is immediately following electrical stimulation, and “post” is after nanosensor fluorescence has returned to baseline. Scale bars represent 10 µm.

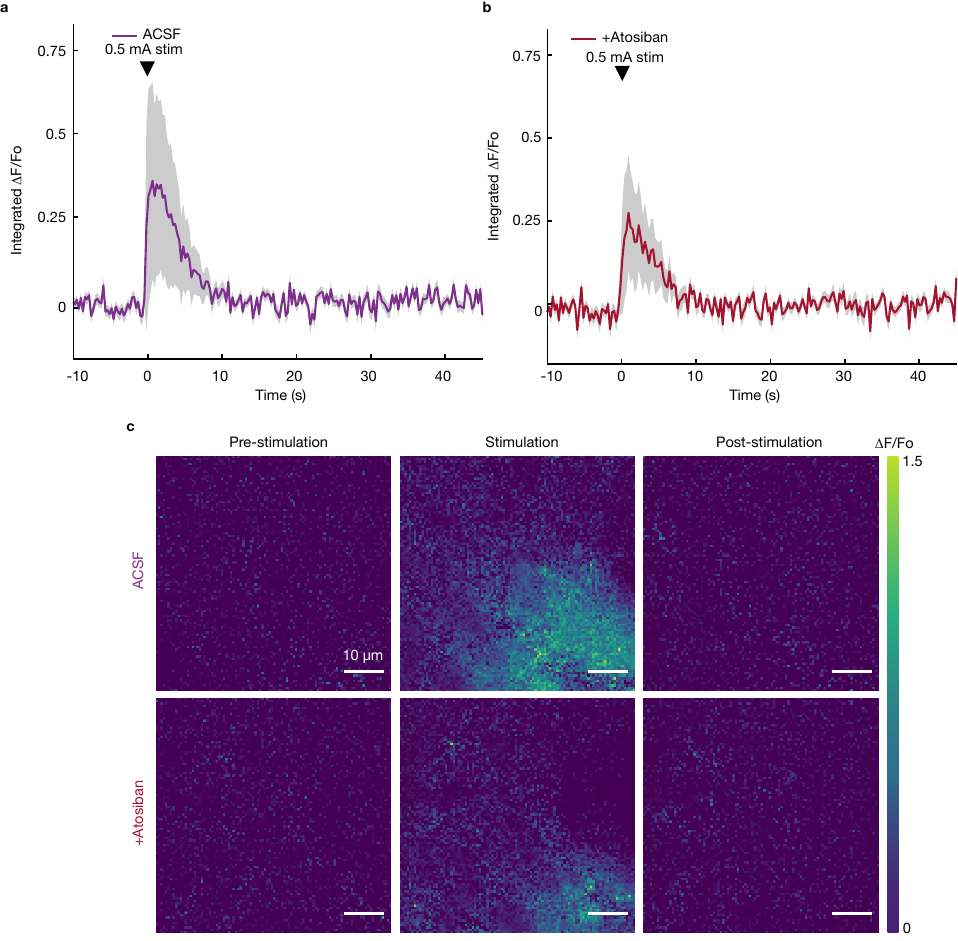

**Supplementary Figure 5 | Analysis of oxytocin release evoked by electrical stimulation and the effect of an oxytocin receptor antagonist in the paraventricular nucleus. a,** *In slice,* mean time trace (purple) and standard deviation (gray) of integrated nanosensor ΔF/F_ο_ for a single 0.5 mA electrical pulse stimulation in the PVN in standard ACSF. **b,** For the same brain slice, mean time trace (red) and standard deviation (gray) of integrated ΔF/F_ο_ for a single 0.5 mA electrical pulse stimulation in 1 µM atosiban in ACSF. **c,** Three images within the same field of view of integrated ΔF/F_ο_ of nanosensor after 0.5 mA electrical stimulation in standard ACSF (top) and 1 µM atosiban in ACSF (bottom). Three frames are represented: “pre” is the baseline before electrical stimulation, “stim” is immediately following electrical stimulation, and “post” is after nanosensor fluorescence has returned to baseline. Scale bars represent 10 µm.

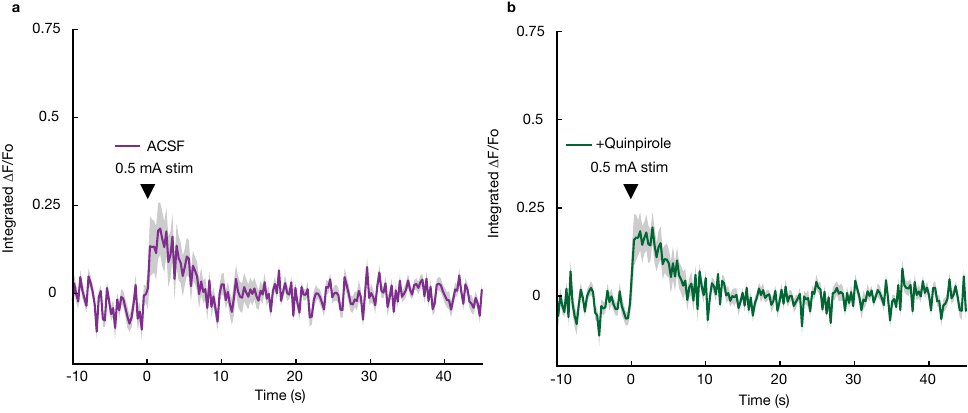

**Supplementary Figure 6 | Analysis of oxytocin release evoked by electrical stimulation and the effect of an D2/D3 receptor agonist in the paraventricular nucleus. a,** *In slice,* mean time trace (purple) and standard deviation (gray) of integrated nanosensor ΔF/F_ο_ for a single 0.5 mA electrical pulse stimulation in PVN in standard ACSF. **b,** Mean time trace (green) and standard deviation (gray) of integrated ΔF/F_ο_ for a single 0.5 mA electrical pulse stimulation in PVN with 1 µM quinpirole in ACSF.

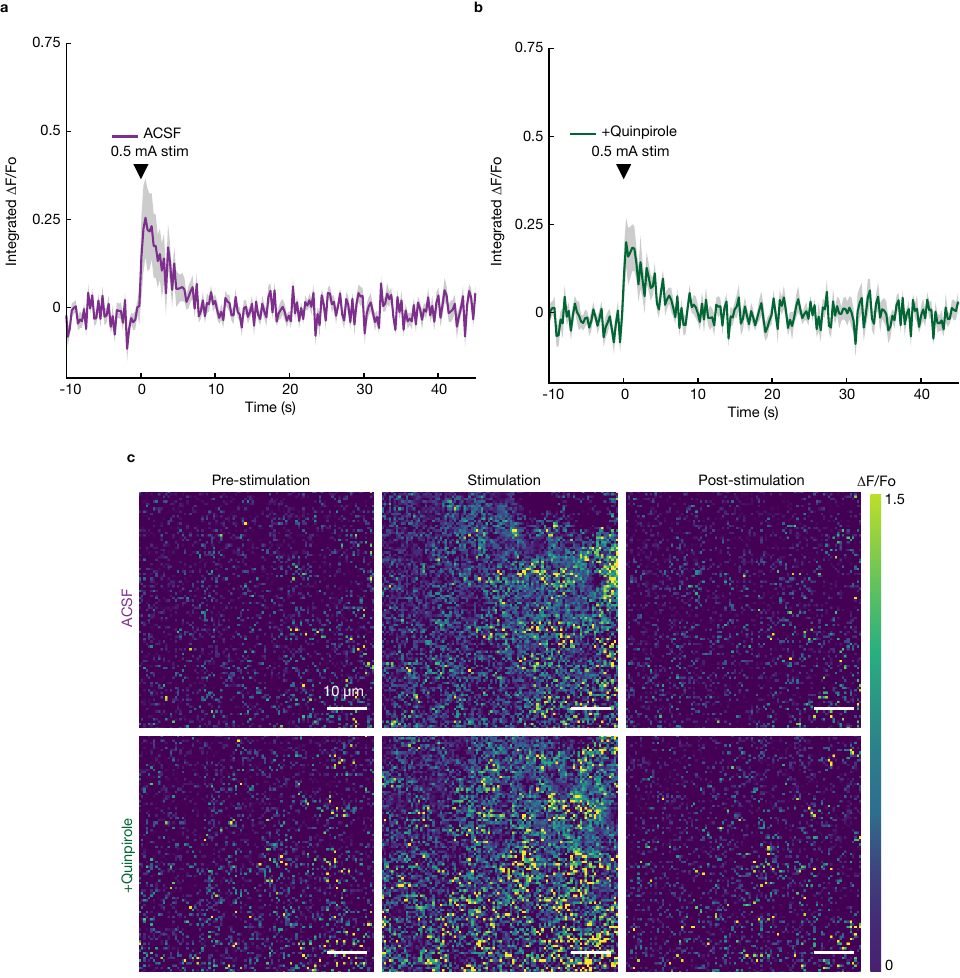

**Supplementary Figure 7 | Analysis of oxytocin release evoked by electrical stimulation and the effect of an D2/D3 receptor agonist in the paraventricular nucleus. a,** *In slice,* mean time trace (purple) and standard deviation (gray) of integrated nanosensor ΔF/F_ο_ for a single 0.5 mA electrical pulse stimulation in PVN in standard ACSF. **b,** Mean time trace (green) and standard deviation (gray) of integrated ΔF/F_ο_ for a single 0.5 mA electrical pulse stimulation in PVN with 1 µM quinpirole in ACSF. **c,** Three images within the same field of view of integrated nanosensor ΔF/F_ο_ following a single pulse 0.5 mA electrical stimulation in standard ACSF (top) and in 1 µM quinpirole in ACSF (bottom). Three frames are represented: “pre” is the baseline before electrical stimulation, “stim” is immediately following electrical stimulation, and “post” is after nanosensor fluorescence has returned to baseline. Scale bars represent 10 µm.

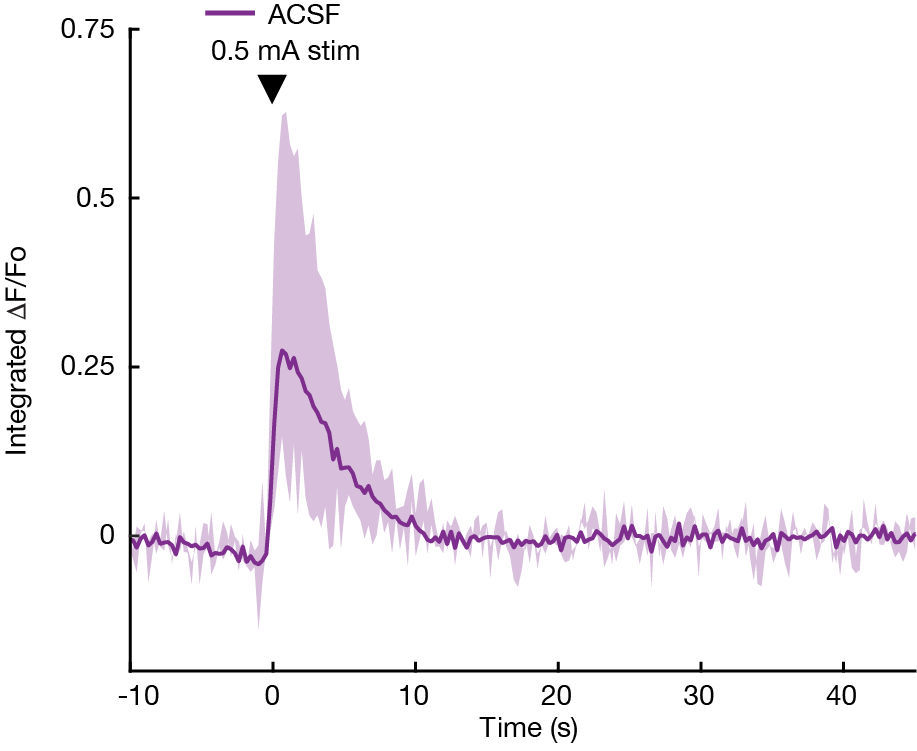

**Supplementary Figure 8 | Analysis of oxytocin release evoked by electrical stimulation.** The range of mean integrated ΔF/F_ο_ traces for twelve 0.5 mA electrical pulse stimulations across 4 biological replicates in standard ACSF are shown (light purple) to demonstrate variation in oxytocin release across stimulations and across acute slices in the PVN. The mean of these traces is shown in dark purple.

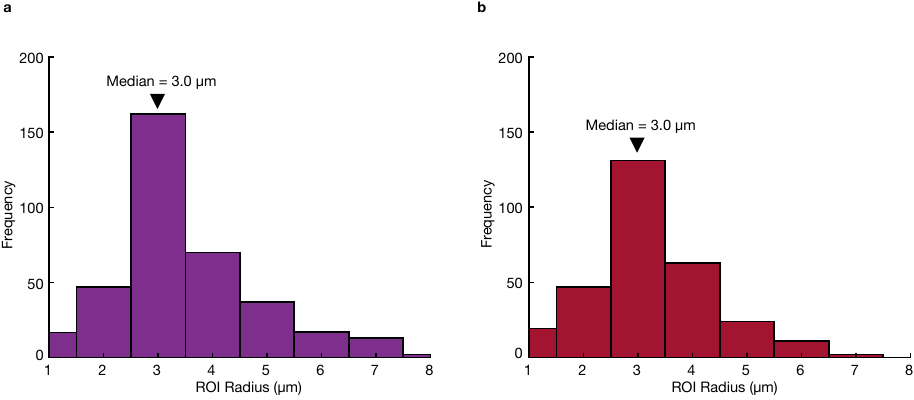

**Supplementary Figure 9 | Region of interest (ROI) analysis of acute slice images to characterize oxytocin release site features. a,** Frequency histogram of ROI sizes across 6 stimulations of the PVN in ACSF. There are 366 total ROIs with a median size of 3 µm. **b,** Frequency histogram of ROI sizes across 6 stimulations of the PVN in 1 µM atosiban in ACSF. There are 297 total ROIs with a median size of 3 µm.
